# Particle Uptake Driven Phagocytosis in Macrophages and Neutrophils Enhances Bacterial Clearance

**DOI:** 10.1101/2021.08.16.456482

**Authors:** Preeti Sharma, Anjali Vijaykumar, Jayashree Vijaya Raghavan, Supriya Rajendra Rananaware, Alakesh Alakesh, Janhavi Bodele, Junaid Ur Rehman, Shivani Shukla, Virta Wagde, Savitha Nadig, Sveta Chakrabarti, Sandhya S. Visweswariah, Dipankar Nandi, Balasubramanian Gopal, Siddharth Jhunjhunwala

## Abstract

Humans are exposed to numerous synthetic foreign particulates in the form of environmental pollutants and diagnostic or therapeutic agents. Specialized immune cells (phagocytes) clear these particulates by phagocytosing and attempting to degrade them. The process of recognition and internalization of the particulates may trigger changes in the function of phagocytes. Some of these changes, especially the ability of a particle-loaded phagocyte to take up and neutralize pathogens, remains poorly studied. Herein, we demonstrate that the uptake of non-stimulatory cargo-free particles enhances the phagocytic ability of monocytes, macrophages and neutrophils. The enhancement in phagocytic ability was independent of particle properties, such as size or the base material constituting the particle. Additionally, we show that the increased phagocytosis was not a result of cellular activation or cellular heterogeneity but was driven by changes in cell membrane fluidity and cellular compliance. A consequence of the enhanced phagocytic activity was that particulate-laden immune cells neutralize *E. coli* faster in culture. Moreover, when administered in mice as a prophylactic, particulates enable faster clearance of *E. coli* and *S. epidermidis*. Together, we demonstrate that the process of uptake induces cellular changes that favor additional phagocytic events. This study provides insights into using non-stimulatory cargo-free particles to engineer immune cell functions for applications involving faster clearance of phagocytosable particulates.

## Introduction

Specialized immune cells utilize the process of phagocytosis for both tissue homeostasis and host defense.^1–3^ As part of host defense, these phagocytic immune cells take up foreign particulates such as microbial pathogens, diagnostic or therapeutic agents,^4^ and micrometer-sized environmental pollutants,^5^ and attempt to degrade them within intracellular compartments. The process of interaction with and uptake of foreign substances may change the phenotype and function of the phagocytic immune cells, a phenomenon widely investigated in the context of microorganisms.^6^ In contrast, the effects of uptake of synthetic particulates on immune cell functions are relatively less explored.

Based on the physicochemical properties of a synthetic particle, reports have suggested that phagocytic immune cells may be activated towards an inflammatory^7–9^ or anti-inflammatory phenotype.^8,10–13^ Additionally, uptake of particles may alter cytokine secretion, chemotaxis behavior, oxidative burst, and nitric oxide generation in these cells.^13–20^ However, it remains unclear if the uptake of particles would affect the ability of a phagocytic immune cell to subsequently phagocytose and neutralize a pathogen.

In this study, we determine how the uptake of particles changes an immune cell’s phagocytic and bactericidal abilities. Using various phagocytic cell types and particles, we demonstrate that uptake of a non-stimulatory cargo-free particle enhances the phagocytic ability of immune cells. We show that this increased phagocytosis is not a result of cellular activation or cellular heterogeneity; instead, the uptake of particles drives subsequent phagocytic events. Finally, we demonstrate that a consequence of the enhanced uptake ability is faster clearance of bacteria both *in vitro* and *in vivo*.

## Results

### Sequential Phagocytosis

Phagocytic immune cells have the ability to engulf multiple particulates. Given the ever-increasing exposure of humans to foreign particulates, phagocytic cells might encounter particulates followed by pathogens in a sequential manner. The question of interest to us was whether the phagocytic ability of these cells is altered after an uptake event. To address this question, we sequentially added particles labeled with two different fluorophores, to cells in culture (Figure 1A). We quantified the fraction of cells that had taken up both the particles (sequential uptake) and compared them to the fraction of cells that had only taken up the particle added second in the sequence (bystander uptake). We observed that sequential uptake was significantly higher than bystander uptake for all combinations of polystyrene (PS) particle (carboxyl-modified surfaces) sizes tested (Figure 1B and supplementary figure 1A). Besides, the number of particles phagocytosed by cells in the second round, quantified as median fluorescence intensity (MFI), was also higher for cells that had taken up particles in the first round (Supplementary figure 1B and 1C). Notably, the number of particles taken up by a cell in the second round significantly increased with the number of particles it phagocytosed in the first round, suggesting dose-dependent priming of phagocyte for the second round of uptake (Supplementary figure 2A and 2B). Furthermore, the phenomenon of enhanced sequential phagocytosis was independent of the time of incubation with particles in the first round (Supplementary 2C), for the timeframes we tested.

**Figure 1:**
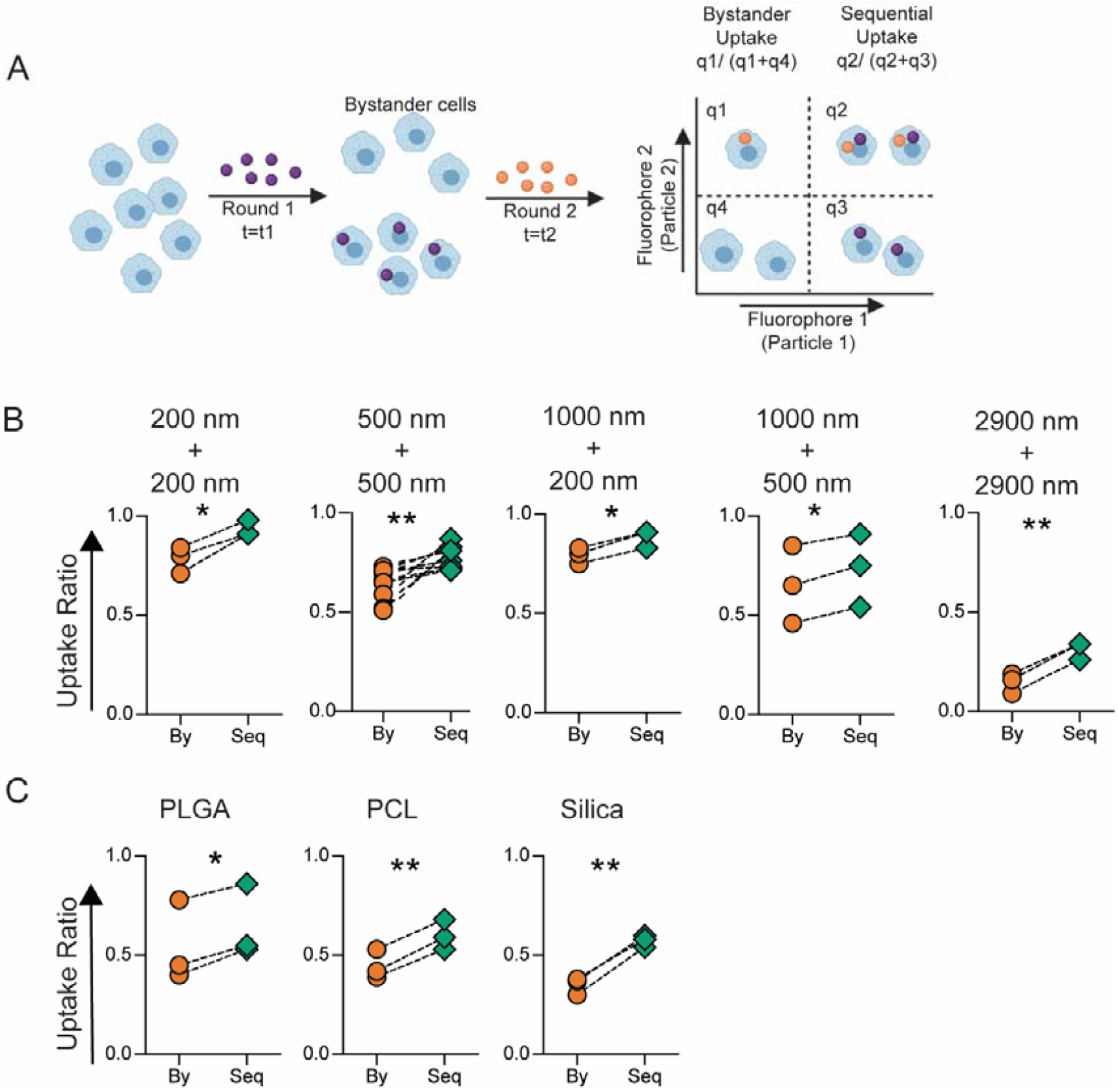
*In Vitro* Sequential Phagocytosis. **A –** Schematic describing the experimental design for calculating bystander and sequential uptake. **B –** Comparison of bystander (By) and sequential (Seq) uptake of carboxylated polystyrene (PS) particles of different diameters by RAW macrophages. In these experiments, t1 and t2 are 2 hours each, and cell to particle ratio was 1:250 for 200 nm, 1:50 for 500 nm , 1:50 for 1000 nm and 1:2 for 2900 nm PS particles. **C** – Sequential uptake with 2-3 µm sized particles composed of PLGA (5 µg/ml), PCL (5 µg/ml) and Silica (1:20) added to RAW macrophages in the first round followed by incubation with 1 µm-carboxylated PS particles (1:10) in the second round of phagocytosis; PLGA = poly (lactic-co-glycolic) acid; PCL = polycaprolactone. Uptake percentages were quantified using flow cytometry. Data sets are representative of *n* ≥ 3 (independent experiments). * = *p* < 0.05 and ** = *p* < 0.01 calculated using paired Student’s *t-*test.

Physicochemical properties of the bulk material and the surface characteristics of particulates are known to dictate the particle-immune cell interaction.^21^ To determine if these properties affected the sequential phagocytosis capacity, we used particles composed of materials approved for clinical use, such as PLGA, PCL and Silica. Uptake of particles made of these materials also resulted in enhanced sequential phagocytosis (Figure 1C). Additionally, cells that take up PS particles whose surfaces do not have a carboxyl group or those that have been modified with polyethylene glycol (PEG) also showed the capacity for increased phagocytosis (Supplementary Figure 3).

The phenomenon of enhanced sequential phagocytosis was observed in another cell line (dHL-60), and among primary monocytes and neutrophils isolated from peripheral venous blood of humans (Figure 2A). Interestingly, this phenomenon was not limited to mammalian systems, as hemocytes isolated from *Drosophila melanogaster* larvae also showed increased sequential phagocytosis (Figure 2B), suggesting that this phenomenon was conserved across phylae. To determine if sequential uptake was observed *in vivo*, fluorescent particles were sequentially administered via the intraperitoneal route in mice, and increased sequential phagocytosis was observed in the peritoneal macrophages (Figure 2C).

**Figure 2:**
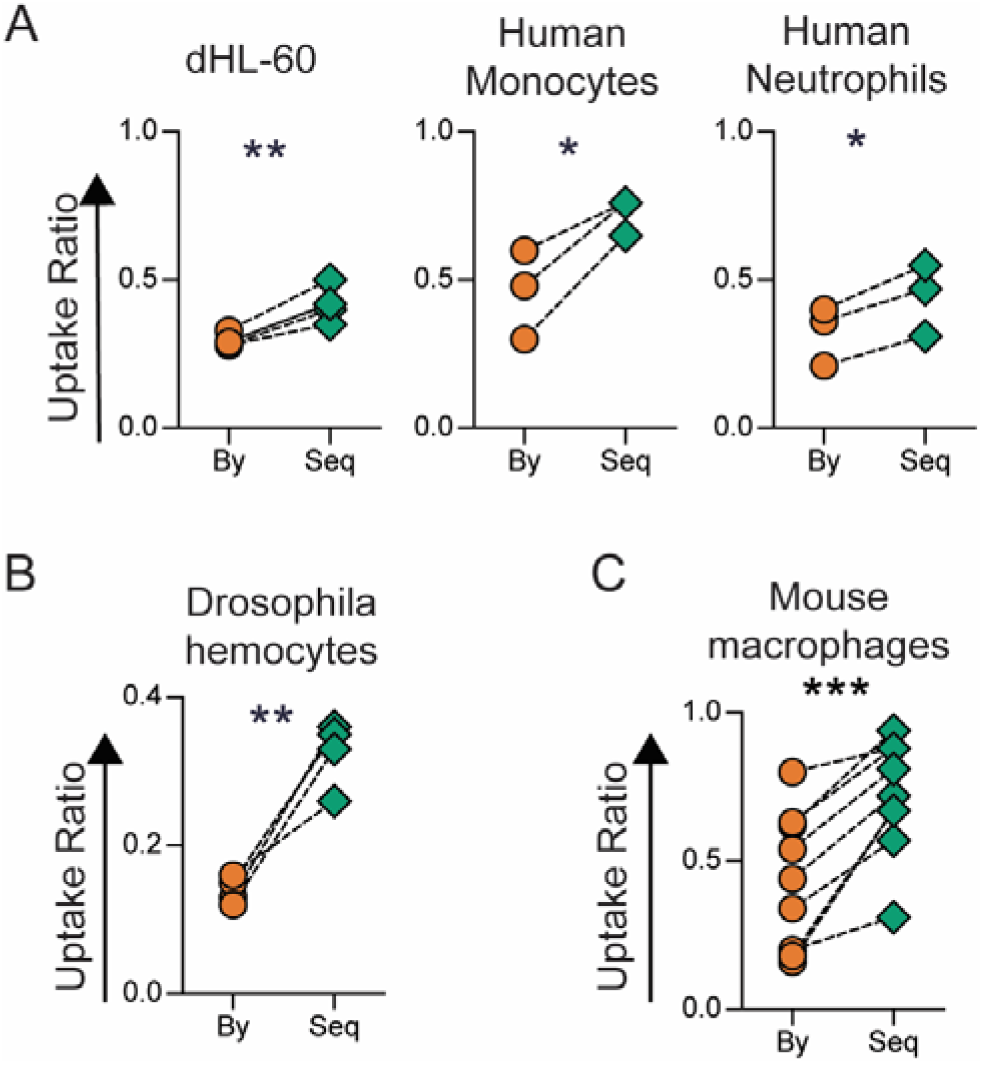
Sequential Phagocytosis among Immune Cells from Humans, Drosophila and in Mice. **A** – Uptake of 500 nm-carboxylated polystyrene (PS) particles by different types of phagocytic immune cells studied using the following cell to particle ratio and incubation times (t1+t2): dHL-60 – 1:100 for 2+2 hours; monocytes – 1:200 for 18+4 hours; neutrophils 1:200 for 4+4 hours. Uptake percentages were quantified using flow cytometry. **B** – *Ex vivo* uptake among hemocytes isolated from Drosophila larvae incubated for 2 hours with 3 µm BSA-TRITC-adsorbed PS particles at a cell to particle ratio of 1:20, followed by incubation with 500 nm-carboxylated PS particles for 2 hours at a cell to particle ratio of 1:200. **C –** Measurement of uptake by mouse peritoneal macrophages (F4/80^+^ cells) following sequential intraperitoneal injection of 6 × 10^6^ 500 nm-carboxylated polystyrene (PS) particles, labelled with different fluorophores, in BALB/c (n=6) or C57BL/6 (n=2) mice. Uptake was quantified using flow cytometry. In all the data sets, “By” indicates uptake by bystander cells and “Seq” represents sequential uptake. Data sets are representative of *n* ≥ 3. * = *p* < 0.05; ** = *p* < 0.01; and *** = *p* < 0.001 calculated using Student’s *t*-test.

We hypothesized that the increase in the ability of cells to phagocytose particles following an initial uptake event could be due to: (i) cellular activation through TLR stimulation during the first round of phagocytosis; or (ii) pre-existing cellular heterogeneity – that is, some cells are inherently more phagocytic; or (iii) the first uptake event drives cellular changes that enhance a cell’s ability to phagocytose substances.

### Cellular activation Does Not Drive Enhanced Sequential Phagocytosis

Phagocytosis of silica and iron oxide particles have been reported to activate immune cells towards pro-inflammatory phenotype characterized by secretion of inflammatory cytokines and generation of reactive oxygen and nitrogen species.^18,20^ Activated immune cells are thought to have an increased phagocytic ability. Further, the presence of contaminating sources of lipopolysaccharide (LPS) on particles could result in increased uptake. We investigated whether the enhanced phagocytic ability observed in our experiments is due to cellular activation following Toll-like receptor (TLR)-4 stimulation. We conducted sequential phagocytosis experiments with RAW cells that were activated with LPS. TLR stimulation with LPS is known to increase the number of cells that will undergo phagocytosis^22^, which was confirmed in the present study as well (Supplementary Figure 4A). We observed enhanced sequential phagocytosis even after uniform pre-activation of cells with LPS (Figure 3A). Next, we blocked the TLR-4 pathway using a chemical inhibitor (TAK-242^23^) prior to the sequential addition of particles. The inhibitor was able to prevent an LPS-mediated increase in overall phagocytosis (Supplementary Figure 4B and 4C). However, even in the presence of this inhibitor, cells showed increased sequential uptake (Figure 3B).

**Figure 3:**
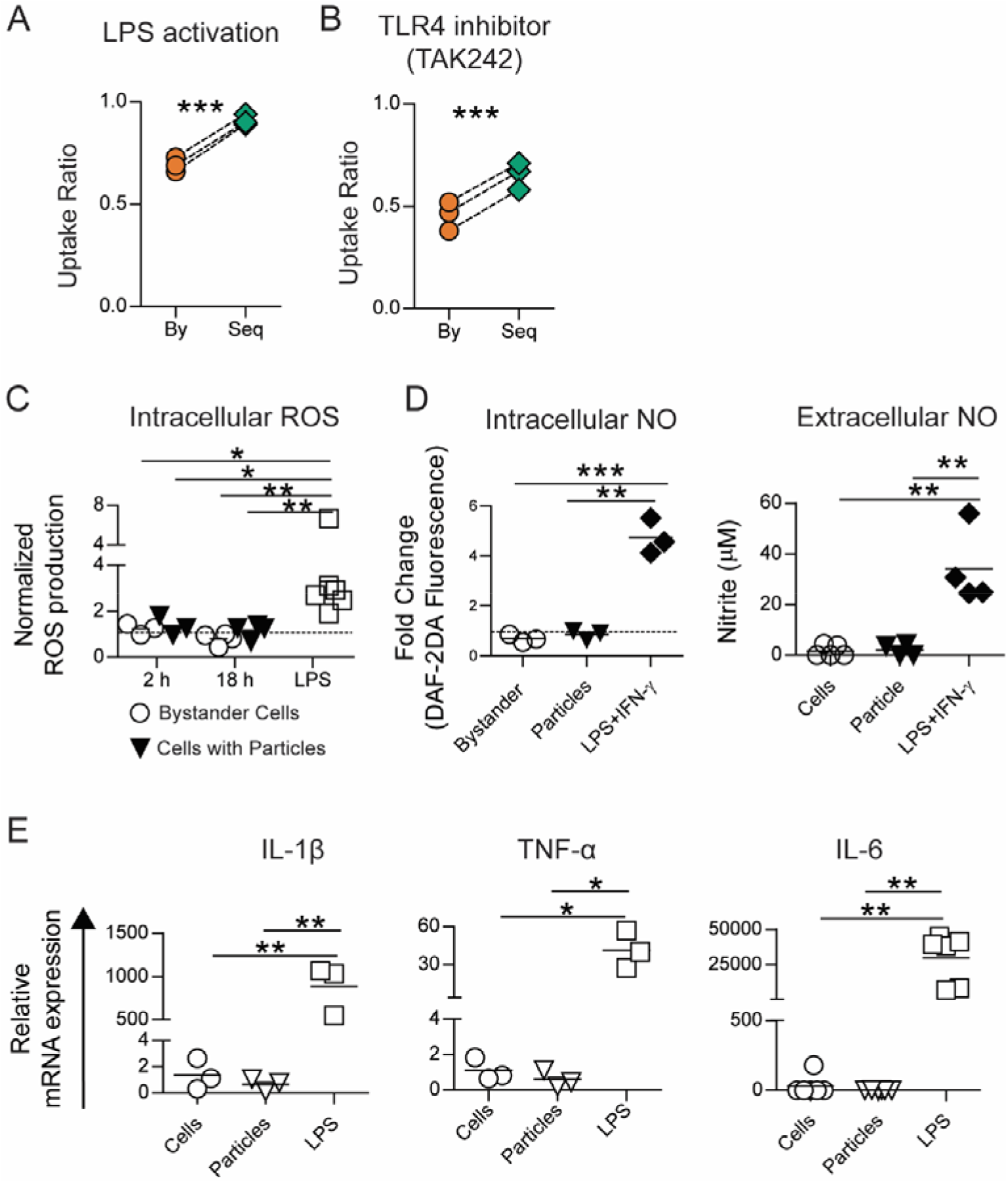
Sequential uptake is not due to TLR based cellular activation. **A** – Uptake of 500 nm-carboxylated polystyrene (PS) particles by RAW macrophages (1:50 cell to particle ratio) following pre-activation of all cells with LPS (1 µg/ml for 18 hours). **B** – Sequential uptake of 500 nm-carboxylated PS particles by RAW macrophages following the treatment of cells with TAK-242 (2 µM for 6 hours) to block TLR-4 mediated cellular activation. **C** – Intracellular reactive oxygen species (ROS) quantified using flow cytometry following incubation of RAW macrophages with 500 nm-carboxylated PS particles (1:50 cell to particle ratio) for 2 or 18 hours, or with LPS for 18 hours. Data are normalized to ROS production in cells that were left untreated for the same times. **D** – Intracellular nitric oxide (NO) production determined after incubating cells with 500 nm-carboxylated PS particles (1:10 cell to particle ratio; lower particle ratio was used due to length of incubation) or LPS and IFN-γ for 36 hours. Extracellular NO production determined after incubating cells with 500 nm-carboxylated PS particles (1:100 cell to particle ratio) or LPS and IFN-γ for 36 hours. **E** – Relative mRNA expression of pro-inflammatory cytokine genes (IL-1β, IL-6 and TNF-α) measured using RT-qPCR. Data are based on *n* ≥ 3 independent experiments (each performed in duplicate). * = *p* < 0.05 ** = *p* < 0.01 and *** = *p* < 0.001 determined using paired Student’s *t*-test or one-way ANOVA.

Activated immune cells are known to produce reactive oxygen species (ROS), nitric oxide (NO) and upregulate the expression of inflammatory cytokines.^24^ We investigated if uptake of PS particles stimulates cells towards an activated phenotype. We observed that while LPS or LPS + IFN□ stimulated cells in culture produced increased levels of intracellular ROS (Figure 3C), NO (Figure 3D) and the inflammatory cytokines IL-1β, IL-6 and TNF-α (Figure 3E), cells that had phagocytosed PS particles did not show any changes in the levels of these molecules compared to controls (naïve cells in culture), showcasing the non-stimulatory nature of these particulates. Further, the anti-inflammatory or M2-type cytokine, IL-10, remained undetectable in all these culture conditions. Collectively, these data suggest that enhanced sequential phagocytosis is not a result of traditional cellular activation caused by TLR stimulation.

### Cellular Heterogeneity is Not Necessary for Enhanced Sequential Phagocytosis

Recent studies ^25,26^ have suggested that differing phagocytic capacities within a given population of immune cells might result in a subset of these cells taking up more bacteria. We used a mouse macrophage cell line to determine if heterogeneity explains the observed increases in sequential uptake. For the first set of experiments, we incubated cells with PS particles at a concentration that resulted in greater than 95% of cells taking up at least one particle, suggesting a close to homogenous population in terms of a phagocytic event. We observe that these cells showed a higher phagocytic ability (percentage of cells with particles) when compared to naïve cells that had not been exposed to particles (Supplementary Figure 5). Separately, we also incubated cells with a lower number of particles that lead to approximately half the cells in a culture-dish taking up particles, resulting in a heterogeneous population in terms of a phagocytic event. These cells were then sorted to separate cells containing particles (particle-positive) from cells that did not take up particles (particle-negative). The sorted cells were then cultured with particles labeled with two different fluorophores in a sequential manner (Figure 4A). The particle-negative cells, which could possibly be considered as a homogenous population of cells with lower phagocytic capacity, also showed enhanced sequential phagocytosis ability (Figure 4B). Additionally, the ‘particle-positive’ cells were not only capable of further phagocytosis but demonstrated increased sequential phagocytosis ability (Figure 4C). These data suggest that increased phagocytic ability is not likely due to cellular heterogeneity alone, at least in this cell line.

**Figure 4:**
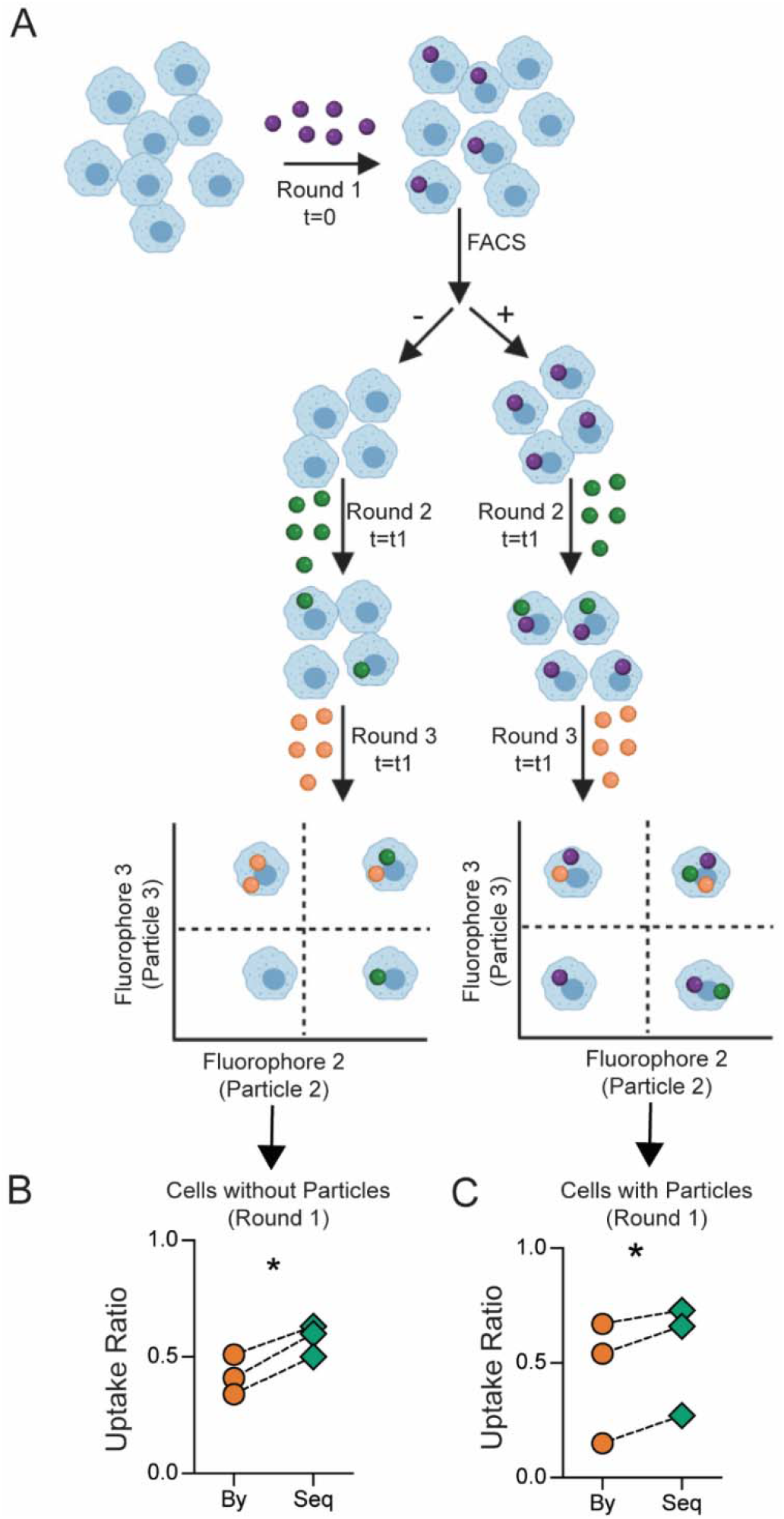
Cellular heterogeneity does not explain sequential uptake in its entirety. **A** – Schematic describing the experimental design of sorting and uptake study. **B** and **C** – Measurement of bystander (By) and sequential (Seq) phagocytosis by cells that did not take up particles prior to sorting (**B**) and those that did take particles prior to sorting (**C**). Following particle types and cell to particle ratio were used: round 1, 500 nm-carboxylated polystyrene (PS-COOH), 1:10; round 2, BSA-TRITC adsorbed 3 µm PS-COOH, 1:5; round 3, 2900 nm PS-COOH, 1:2. FACS stands for fluorescence-activated cell sorting. Data are based on *n* = 3 independent experiments (each performed in duplicate). * indicates *p* < 0.05 determined using paired Student’s *t*-test.

### Phagocytosis Induces Changes in Membrane Fluidity and Cellular Stiffness

Having observed that neither cellular activation nor cellular heterogeneity could explain the enhanced sequential phagocytosis phenomena, we explored if phagocytosis induces changes in cells that might result in increased uptake. Both the cell membrane and the cytoskeleton are actively involved in phagocytosis, and hence we assessed changes to these structures following uptake of PS particles. Using a fluorescent probe that detects perturbations in membrane phase properties, we determined that the fluidity (measured based on generalized polarization value of the dye) of the membranes of cells that have phagocytosed PS particles was significantly higher as compared to naïve cells (Figure 5A). Methyl-beta-cyclodextrin (MβCD), a cholesterol-depleting agent, was used as a positive control in these experiments and shows the highest increases in membrane fluidity. Increased membrane fluidity has been associated with higher phagocytic ability^27^, and the above observation suggests that one possible explanation for the enhanced sequential phagocytosis could be changes in fluidity of cell membranes.

**Figure 5:**
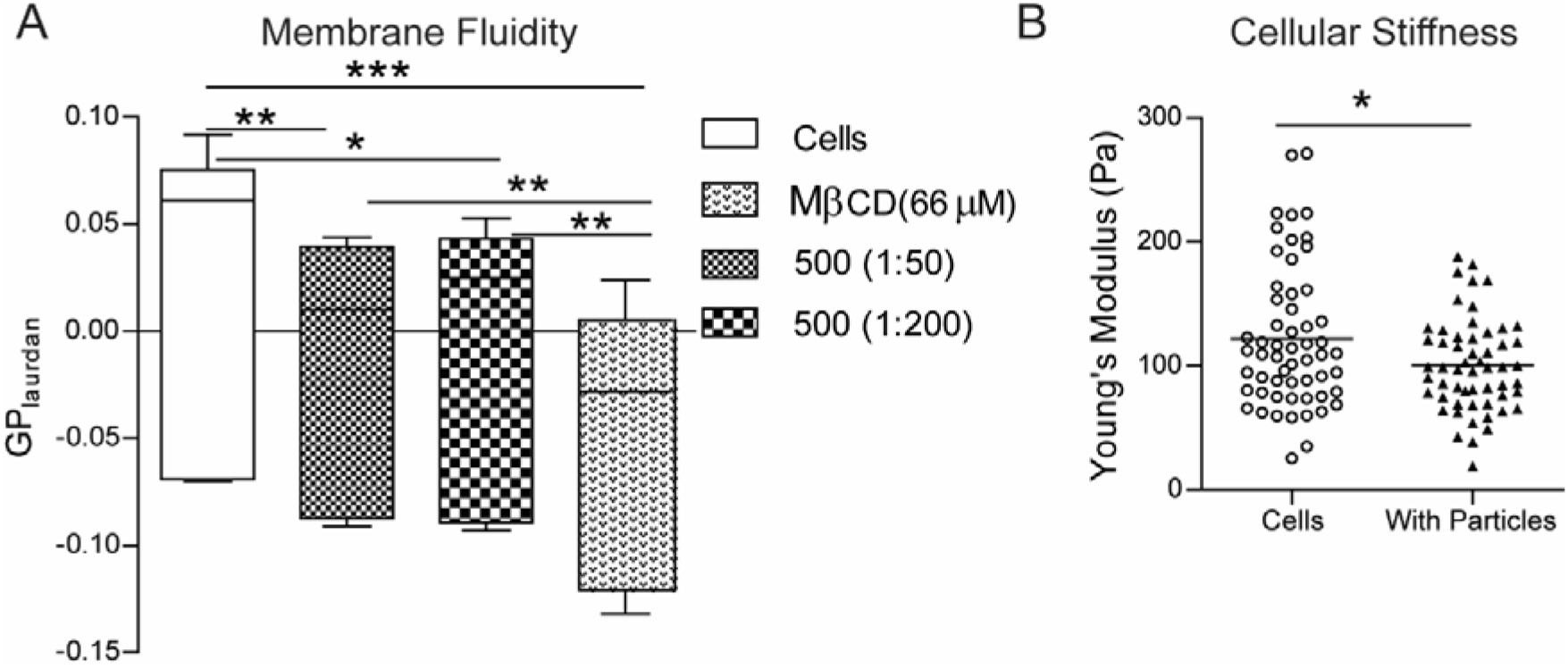
Phagocytosis induced changes in macrophage membrane fluidity and cellular stiffness. **A** – Measurement of membrane fluidity using the generalized polarization (GP) value for the membrane dye Laurdan in RAW macrophages that were incubated with 500 nm-carboxylated polystyrene (PS) particles at two different cell-to-particle ratios and compared to the negative control of naïve cells (cells) as well as the positive control of cells treated with methyl-beta-cyclodextrin (MβCD). *n* = 6 independent experiments. * indicates *p* < 0.05, ** indicates *p* < 0.01 ** and *** indicates *p* < 0.001 calculated using one-way ANOVA followed by Bonferroni post-hoc test for comparison of multiple groups. **B** – Measurement of apparent modulus of a cell using atomic force microscopy. Young’s modulus of naïve cells (Cells) and cells that have been treated with 500 nm-carboxylated PS particles (With Particles) is plotted, with each dot representing a single cell. Data sets are representative of at least 60 cells measured across 3 independent experiments. * indicates *p* < 0.05 calculated using unpaired *t*-test with Welch’s correction.

Additionally, a cell must change its shape to internalize phagocytic targets. Hence, we measured the ability of a cell to deform by estimating the cellular stiffness using force-distance spectroscopy in an atomic force microscope. The Young’s modulus of cells that had phagocytosed 500 nm-carboxylated PS particles was significantly lower than naïve cells (Figure 5B and Supplementary Figure 6), indicating an enhanced ability to deform, which might partly explain the increase in phagocytic capacity.

### Increased *in vitro* clearance of *E. coli*

The primary function of phagocytic immune cells is to recognize, phagocytose and degrade pathogenic microorganisms.^3,28^ Phagocytic cells that have internalized particles might encounter bacteria; hence, we examined the effect of particle uptake on the internalization and killing of bacteria by phagocytic immune cells. We used *E. coli* as a model system for these studies. The bacteria were added to macrophage cell-line cultures (MOI 3.3 - 44) that had either been exposed to PS particles (at a dose that resulted in >90% cells having particles) or naïve cells. In concurrence with our enhanced sequential uptake data, at 2 hours post *in vitro* infection, we observed increased internalization of *E. coli* by RAW macrophages that had taken up particles compared to naïve cells (Figure 6A). Higher bacterial numbers (measured as colony-forming units (CFUs)) were observed in cells with particles at 6 hours too; however, by 18 hours, the bacterial numbers in cells with particles were similar to that of naïve cells. Upon calculation of the clearance of bacteria, which is the number internalized initially to the number that remained at the end of measurements, we observe that cells with particles had cleared a greater percentage of bacteria (Figure 6B).

**Figure 6:**
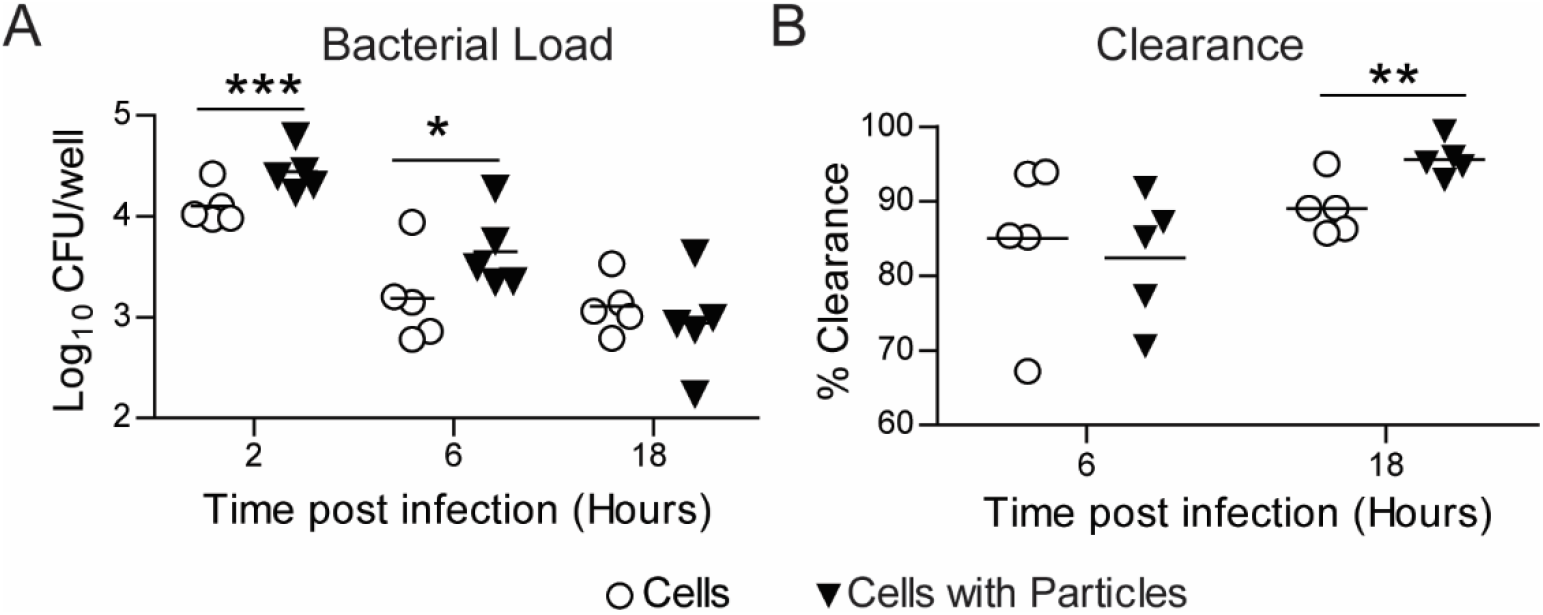
*E. coli* Clearance - *in vitro*. **A –** Measurement of *E. coli* numbers inside RAW macrophages at different times following exposure of cells to bacteria. Cells were previously treated with 500 nm-carboxylated polystyrene particles for 18 hours (Cells with Particles), and naïve cells (Cells) were used as controls. Enumeration of intracellular bacteria as colony forming units (CFU) obtained by plating cell lysate at specified time points. **B –** Change in bacterial load measured over time as percentage intracellular bacteria killed by time ‘t’ compared to 2-hour time-point. Data sets are representative of *n* = 5 independent experiments. * = *p* < 0.05, ** = *p* < 0.01 and *** = *p* < 0.001 calculated using Student’s *t*-test.

### Unaltered invasion and replication of *S*. Typhimurium inside macrophages with particles

Bacteria such as *Salmonella* are known to invade, survive and replicate inside phagocytic immune cells actively. So, we evaluated if the phenomenon of enhanced sequential phagocytosis would alter the interaction of such bacteria with phagocytes. We added *S*. Typhimurium, an intracellular pathogen, to RAW macrophage cultures (*in vitro*) (MOI 5 - 50) and observed that the numbers of intracellular bacteria were not different in cells with PS particles when compared to cells without particles (Supplementary Figure 7). This observation is not surprising as these specific bacteria actively invade macrophages, in addition to being taken up by phagocytosis. Further, the numbers of these bacteria continued to remain equally high in both cells with particles and naïve cells, which might also be expected as we show that the uptake of the non-stimulatory cargo-free PS particle only affects the cell’s phagocytic ability and not the cell’s activation and killing mechanisms such as ROS, NO or cytokine production.

### Phagocytosis of non-stimulatory particles increases the rate of *E. coli* clearance *in vivo*

Next, we investigated the effect of particle phagocytosis by immune cells on the bacterial killing kinetics using an animal model of bacterial infection. For this, particles were injected in C57BL/6 mice via intraperitoneal route 2 hours prior to *E. coli* infection at the same site. Bacterial load was measured in the peritoneal exudate, within peritoneal exudate cells, and in distant sites such as the spleen, kidney and liver. Mice injected with particles showed significantly lower bacterial load in peritoneal exudate cells (intracellular) (Figure 7A) and peritoneal exudate (Figure 7B) at 6 hours after bacterial injection compared to mice injected with saline followed by bacteria (controls). Additionally, the bacterial burden was found to be significantly reduced in the spleen (Figure 7C) and lowered but not statistically significant in the kidney (Figure 7D) of mice injected with particles. In all the mice, we did not detect any bacteria infiltrating the liver.

**Figure 7:**
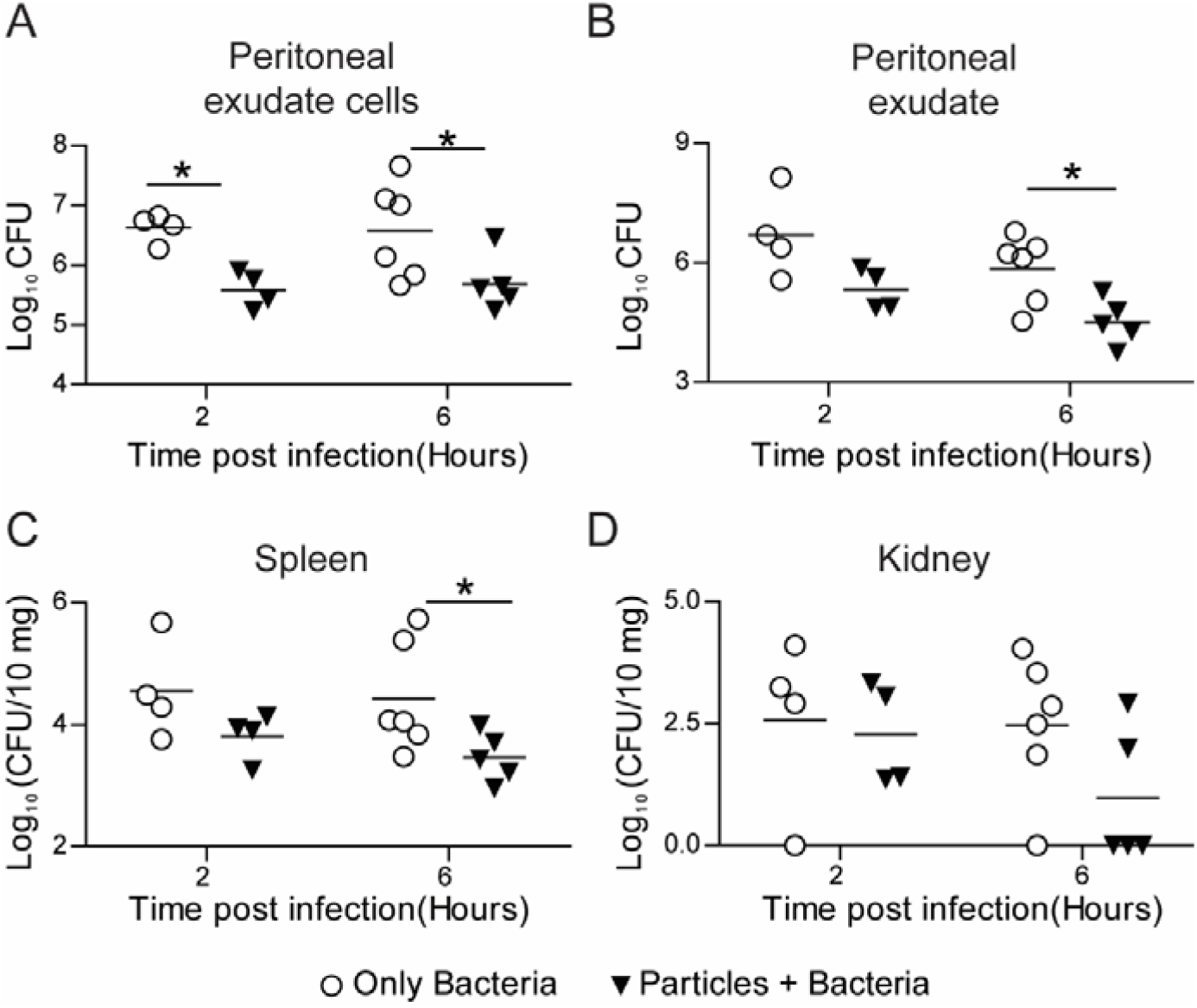
*E. coli* Clearance - *in vivo*. Kinetics of clearance of *E. coli* from C57BL/6 mice following intraperitoneal injection of 10^10^ 500 nm-carboxylated PS particles (or saline), and subsequent injection (2 hours later) of 5 × 10^7^ bacteria per mouse. Bacterial load measured as CFU from lysates of **A –** peritoneal exudate cells, **B –** peritoneal exudate, **C –** spleen and **D –** kidney are plotted against time post bacterial injection (organ CFU plotted as per 10 mg of organ weight) . Each point on the plot corresponds to one mouse. Mann-Whitney test was performed for statistical comparison. * indicates *p* ≤ 0.5.

The total number of immune cells in the peritoneal exudate 2 hours after particle injection was not significantly different from the saline-injected mice, indicating that the faster clearance of bacteria was not due to increased immune cell infiltration (Supplementary Figure 8). The faster killing of bacteria could also occur if immune cells are activated in mice prior to administration of bacteria. Thus, to determine the activation status of immune cells after they have taken up particles, we compared the expression of various activation markers on isolated mouse peritoneal macrophages, which had phagocytosed particles, to that of untreated naïve peritoneal macrophages from the same animals. There was no significant difference in the expression of CD11b, CD38, CD54, CD62L and CD86 between naïve cells and cells with particles (Supplementary Figure 9). Together, these data suggest prophylactic particle administration, which results in enhancing the phagocytic capacity of phagocytes, leads to faster clearance of *E. coli*.

### Phagocytosis of non-stimulatory particles increases the rate of *S. epidermidis* clearance *in vivo*

As *E. coli* is killed by most phagocytic immune cells, we next chose to use a pathogen that is primarily killed by a specific phagocytic immune cell. *S. epidermidis*, a nosocomial pathogen associated with heavy clinical burdens, is primarily neutralized by neutrophils.^29^ To investigate if enhancing the phagocytic ability of immune cells can effectively contain the spread of *S. epidermidis*, we injected particles and bacteria sequentially. Bystander and sequential uptake were determined in neutrophils, macrophages, and monocytes from the peritoneal cavity (Supplementary Figure 10). In concurrence with *in vitro* data on enhanced sequential phagocytosis after particle uptake, our *in vivo* data shows that cells containing particles phagocytose a higher number of bacteria than cells without particles in the same mouse or compared to cells in mice that were not injected with particles (particle-naïve mouse) (Figure 8A). Particle-injected mice showed a trend of lower bacterial burden in the peritoneal space (Figure 8B and 8C). Importantly, in particle-injected mice, we observed that the dissemination of bacteria to distant organs was significantly lowered, with a complete absence of bacteria in the kidney (Figure 8D) and close to zero bacteria after 24 hours in the spleen (Figure 8E), as compared to particle-naïve mice. These data also indicate that prophylactic administration of non-stimulatory particles results in faster clearance and prevention of bacterial spread.

**Figure 8:**
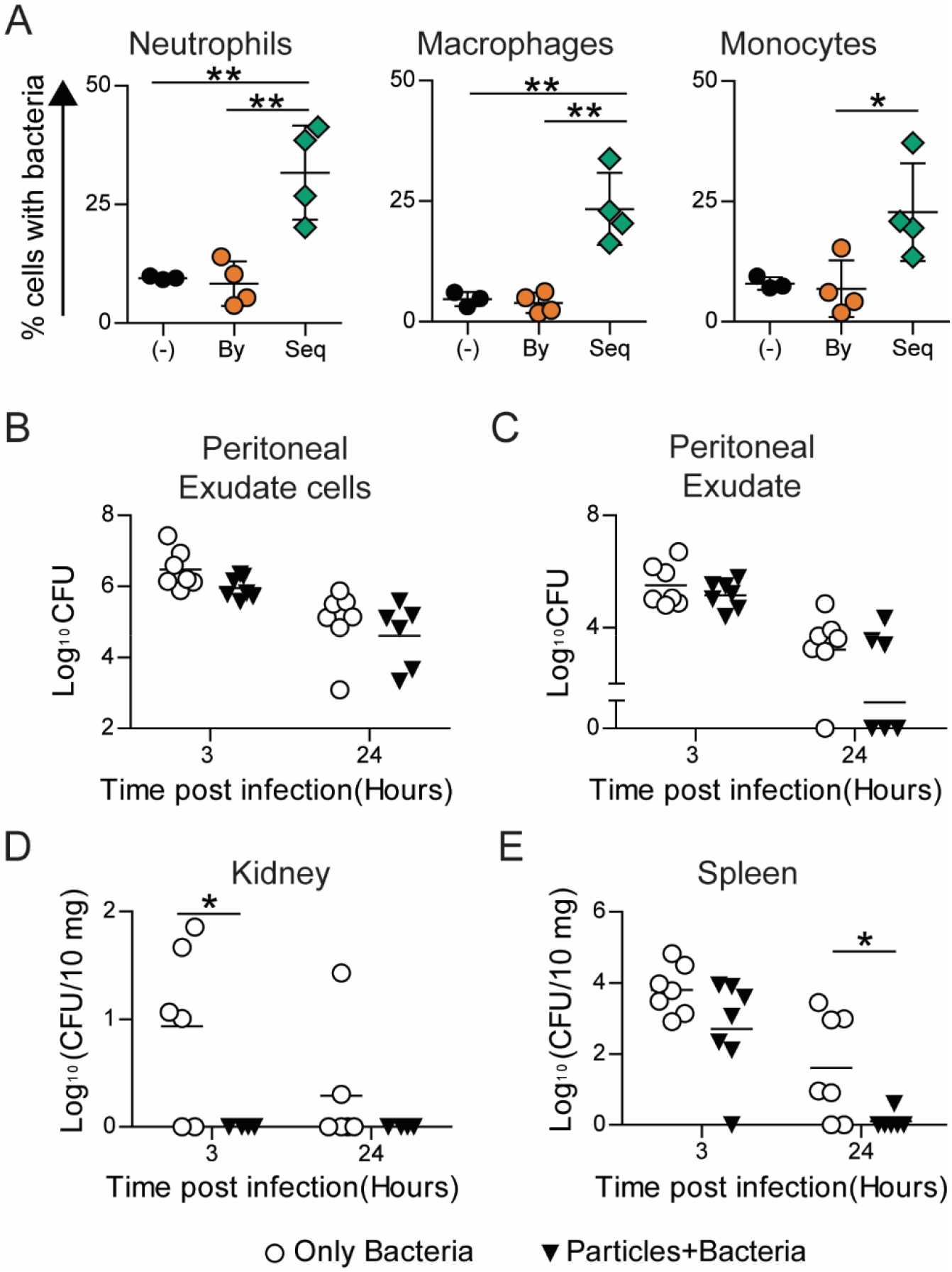
*S. epidermidis* Clearance. **A** – Uptake of fluorescent 500 nm-carboxylated polystyrene (PS) particles and GFP-expressing *S. epidermidis* following their *in vivo* intraperitoneal administration (in C57BL/6) in a sequential manner. (-) indicates mouse injected with saline followed by bacteria, “By” indicates bystander cells and “Seq” indicates sequential uptake cells in mice injected with particles followed by bacteria. **B-E** – Kinetics of bacterial clearance determined in C57BL/6 mice following intraperitoneal injection of 10^9^ 500nm-carboxylated PS particles or saline, and subsequent injection of ∼3 × 10^8^ *S. epidermidis*. Bacterial load measured as colony forming units (CFU) from lysates of **B** – peritoneal exudate cells, **C** – peritoneal exudate, **D** – kidney and **E** – spleen (plotted as CFU per 10 mg of organ weight) are plotted against time post infection. Each data point corresponds to one mouse. Mann Whitney test was performed for statistical comparison. * indicates *p* ≤ 0.5.

## Discussion

*In vivo* administration of particles results in a large proportion of them being sequestered by phagocytic immune cells. A recent meta-analysis of nanoparticle-based cancer therapeutics by Warren Chan and colleagues showed that only 0.7% of the administered particles reach the target tumor site in mouse models^30^, implying that most of the particles are sequestered by phagocytic cells. Even if the particles are degradable, once phagocytosed, they may be associated with the immune cells for days to weeks.^31,32^ Hence, apart from understanding *in cellulo* and *in vivo* fate of the particulates, which is well studied^33–36^, it is crucial to understand the impact of particulate phagocytosis on the immune cell’s functionality.

A cell’s capacity to phagocytose particulates depends on factors such as its polarization state, expression levels of various proteins associated with phagocytosis, and environmental cues such as cytokines present in the extracellular milieu.^37,38^ A change in any of these factors may be used to modulate its uptake capacity. For example, stimulating macrophages with LPS or IFN□, which activate NF-kB, results in heightened phagocytic activity.^22^ However, these methods to modulate phagocytic activity are generally accompanied by other cellular changes, such as the production of inflammatory cytokines and reactive oxygen and nitrogen species by M1 phenotypic cells, which is likely to cause damage to surrounding cells and tissues.^39^ Contrasting these conventional methods of cellular activation, we show that monocytes, macrophages and neutrophils may be driven towards a phenotype that shows enhanced phagocytic activity without eliciting an inflammatory immune response. This observation is independent of particle size, is conserved across various phagocytic cell types and is seen in different organisms, suggestive of its universality.

Phenotypic and functional heterogeneity among immune cells is now well recognized.^40^ Hellebrekers et al. showed that neutrophils have increased phagocytic capacity following *S. aureus* uptake^25^, while Sachdeva et al.^26^ showed that endocytosis increases phagocytic capacity in macrophage cell lines. Both these studies attributed the increased phagocytosis to heterogeneity among cells, with a unique highly endocytic/phagocytic cell population responsible for the increased uptake. While heterogeneity in phagocytic capacity exists, our cell-sorting data imply that an increase in phagocytic capacity may also be induced. Cells that did not initially take up particles (bystander cells) show enhanced sequential phagocytosis when provided with new phagocytic targets in our cell-sorting experiment, suggesting that uptake is able to drive additional phagocytosis. While heterogeneous cells may exist even after cell sorting, we are unlikely to expect a large proportion of them to be highly phagocytic cells. Additional proof for uptake-driven increases in phagocytosis comes from the changes we observe to the fluidity of cell membranes and the Young’s modulus of a cell.

Modulation of phagocytic immune cell function may prove beneficial for a wide range of diseases. Generally, such modulation is achieved through therapeutic molecules that may be delivered with the help of particles. However, a few recent studies have shown that non-stimulatory cargo-free particles may also reprogram immune cell behavior and have demonstrated their use in the treatment of conditions such as inflammatory bowel disease, acute lung injury, sepsis and West Nile infection^41–44^. However, the effects of particles observed in these studies is thought to be due to the diversion of immune cells away from the inflammatory site to secondary lymphoid organs. Highlighting the modulation at the cellular level, our results suggest that phagocytosable cargo-free non-stimulatory particulates may be used in conditions that are marked by impaired phagocytosis. We demonstrate that one such application is the increased rate of clearance and reduction in the systemic dissemination of *E. coli* and *S. epidermidis*. Other possible applications could include assisting with clearance of pathogens from lungs in patients with impaired phagocytic activity^45,46^, faster clearance of apoptotic cells in inflammatory diseases^47,48^ and in improving wound-healing.

Finally, it is important to note that the non-stimulatory cargo-free particulates change an immune cell’s phagocytic ability but not killing capacity. So, the strategy of using particulates for faster bacterial clearance is unlikely to work for pathogens that have active evasion or survival mechanism. Also, the increase in phagocytic ability is brought about after particle uptake, and hence the particle administration may only be used for prophylactic treatments and not as a therapeutic.

In conclusion, we show that uptake of particles drives additional phagocytosis in immune cells. Hence, we may utilize non-stimulatory cargo-free particles to increase the phagocytic ability of immune cells that could have applications in improving bacterial clearance. The work presented here also indicates that particles may have unintended effects on the functionality of immune cells, which should be addressed when evaluating the compatibility of newly designed nano- and micro-particulates for clinical use.

## Materials and methods

### *In vitro* sequential phagocytosis

RAW 264.7 cells were cultured in DMEM (Lonza, India) supplemented with 10% FBS and 1% antibiotic-antimycotic solution (Thermo-fisher Scientific, USA). HL-60 cells were cultured in IMDM (Merck, USA) supplemented with 20% FBS and 1% antibiotic-antimycotic solution. These cells were induced towards granulocytes (d-HL60) with 1.25% DMSO (Merck), for 3 days. Both cell lines were seeded at a density of 2 × 10^5^ cells per well (of a 24-well plate) and incubated for 2 hours at 37°C under 5% CO_2_. For phagocytosis studies, carboxylated polystyrene (PS) particles of various sizes (200 nm, 500 nm, 1000 nm and 2900 nm) labeled with different fluorophores (Dragon-green or Flash-red) were used (Bangs Laboratories, Indiana, USA). Fluorescent bovine serum albumin (BSA) loaded ∼2500 nm-sized Poly(lactic-co-glycolic acid) (PLGA), and ∼2000 nm-sized polycaprolactone (PCL) particles were synthesized using the double emulsion solvent evaporation method.^49^ Silica particles with a diameter of 3000 nm (Bangs Laboratories) were tagged for fluorescence by adsorbing fluorescently labelled BSA on the particle surface. For the first round of phagocytosis, particles labeled with fluorophore 1 were added to the cells at different ratios (exact ratios specified in the figure legends). After co-incubation of cells and particles for various time intervals, media containing free-floating particles was removed, cells were washed with 1 X PBS to ensure minimal to no residual particles, and fresh media was added. Cells were then incubated with particles labelled with fluorophore 2 for a specified time interval. Finally, media was removed, wells washed with 1 X PBS, cells scraped, stained with 2 µg/ml propidium iodide (PI) solution, and analyzed on a flow cytometer.

For experiments involving lipopolysaccharide (LPS) induced cellular activation, RAW cells were treated with 1 µg/ml of LPS (Merck) for 12-18 hours prior to particle addition. Inhibition of the TLR-4 pathway was achieved by treating cells with 2 µM TAK-242 (Merck) for 6 hours before adding particles. For studies involving cell sorting, RAW cells were seeded in a T-25 flask, incubated with fluorescent PS particles for 2 hours and sorted using BD FACSAria™ Fusion (Becton Dickinson, USA) equipped with a 488 laser, under a two-way purity sort setting. Post sorting, particle-positive and particle-negative cells were separately cultured in 24-well tissue culture plates and allowed to adhere for 2 hours, after which a sequential phagocytosis experiment was performed as described above for RAW cells.

### Flow cytometry

Experiments involving flow cytometry were performed on BD FACSCelesta™ (Becton Dickinson) and analyzed using FlowJo (Tree Star, Ashland, OR, USA).

### *Ex vivo* sequential phagocytosis

Studies involving human blood were approved by the Institutional Human Ethics Committee at the Indian Institute of Science (IISc) (approval number 5-15032017). For the *ex vivo* sequential phagocytosis experiment, venous blood was collected in EDTA coated tubes, from healthy volunteers, after obtaining informed consent. PBMCs were isolated using histopaque (Sigma Aldrich, USA) density gradient centrifugation. In brief, 5 ml of whole blood was carefully laid over 7.5 ml of histopaque and centrifuged at 500 rcf for 20 min (with the brake turned off) at room temperature. The PBMC layer was collected, and cells were counted, seeded and allowed to adhere in 24-well cell culture plate for one hour at 37°C under 5% CO_2_. The bottom-most layer of the density gradient, containing neutrophils and RBCs, was collected separately, and RBCs were lysed using Ammonium-Chloride-Potassium (ACK) lysis buffer (10 ml lysis buffer per 1 ml blood for 8 min at room temperature). The lysis reaction was quenched with 10 volumes of 1 X PBS. The tube was further centrifuged at 400 rcf for 5 min, the supernatant discarded, and the neutrophil enriched pellet resuspended in DMEM cell culture media. Neutrophils and PBMCs were counted and seeded in 24-well plates at a density of 1 × 10^5^ cells per well. Sequential phagocytosis experiments were performed as carried out for RAW cells.

### Sequential uptake studies in Drosophila hemocytes

Transgenic *hml*Δ*Gal4, UAS-GFP Drosophila* stocks (Bloomington stock number 30142) were maintained at 25°C on a 12 h light/ 12 h dark-cycle in a medium comprising 8% cornmeal, 4% sucrose, 2% dextrose, 1.5% yeast extract, 0.8% agar supplemented with 0.4% propionic acid, 0.06% orthophosphoric acid and 0.07% benzoic acid. Twenty 3^rd^ instar larvae were dissected, allowed to bleed into buffer consisting of Ringer’s solution and 1 mM phenylthiourea (PTU) for 30 seconds, and passed through a 100 µm mesh to collect single-cell suspension. Sequential uptake experiments were performed on these cells by incubating them with 3 µm BSA-TRITC adsorbed PS particles (cell to particle ratio of 1:20) for 2 hours at room temperature, removing non-phagocytosed particles by centrifugation followed by the addition of 500 nm-carboxylated PS particles (cell to particle ratio of 1:200) for 2 hours. Finally, the cells were washed, and hemocytes (identified based on GFP expression) were analyzed on a flow cytometer for particle uptake.

### *In vivo* sequential phagocytosis

All animal studies were conducted in accordance with the Control and Supervision Rules, 1998 of the Ministry of Environment and Forest Act (Government of India), and the Institutional Animal Ethics Committee, IISc. Experiments were approved by the Committee for Purpose and Control and Supervision of Experiments on Animals (permit numbers CAF/ethics/546/2017 and CAF/ethics/718/2019). Animals were procured from either IISc’s breeding facility (BALB/c mice) or from Hylasco Bio-Technology Pvt Ltd. (a Charles River Laboratories Subsidiary, for C57BL/6 mice). BALB/c or C57BL/6 mice (8–14-week-old) were injected intraperitoneally with 6 × 10^6^ 500 nm-carboxylated fluorescent PS particles. After 2 hours, mice were injected via the intraperitoneal route with 500 nm-carboxylated PS particles labelled with a different fluorophore. Mice were euthanized after 2 hours, and peritoneal exudate was collected by performing peritoneal lavage and cells stained with Ly6G (clone 1A8) and F4/80 (T45-2342) (BD Biosciences) for 20 min at 4 °C. Finally, the cells were stained with PI before being run on the flow cytometer to determine bystander and sequential uptake.

### Reactive Oxygen Species (ROS) assay

ROS produced by immune cells after phagocytosis of PS particles was assessed using an intracellular probe, Dihydrorhodamine (DHR) 123 (Merck). After the incubation of cells with particles for 2 and 18 hours, media was removed, and cells were incubated in fresh media containing 5 µM of DHR for 15 min at 37°C. Following this, the probe was quenched with excess 1 X PBS, cells washed and scraped, stained with PI, and median fluorescence intensity of cells determined using a flow cytometer. For positive control for ROS production, RAW cells were stimulated with LPS (1 µg/ml) for 12-18 hours.

### Nitric Oxide (NO) assay

For extracellular NO measurement, RAW cells were seeded at a density of 1 × 10^5^ cells per well in a 24-well plate and incubated with PS particles for 36 hours. To determine extracellular NO production, 50 µl aliquot of cell culture supernatant was mixed with 150 µl of Griess reagent in a 96-well plate. The amount of NO_2_^-^ was determined from a standard curve of Sodium Nitrite. As a positive control for NO production, cells were stimulated with a combination of LPS (100 ng/ml) and IFN□ (100U/ml) (PeproTech, Israel) for 36-hours.

For intracellular NO measurement, RAW cells were seeded at a density of 1 × 10^5^ cells per well in a 24-well plate. Cells were incubated with PS particles for 36 hours, following which, media was removed and, cells were incubated with 5 µM 4,5-diaminofluorescein (DAF-2A, Sigma-Aldrich, USA) for 15 min at 37 °C in dark. The fluorescent probe was quenched with 1 X PBS, cells were scraped, stained with PI, and fluorescence intensity (corresponding to the amount of NO present), was measured on a flow cytometer using excitation and an emission wavelength of 488 and 520 nm, respectively.

### Real-time quantitative PCR (RT-qPCR) Assays

RAW cells, 4 × 10^5^ per well, were seeded in a 24-well plate and allowed to adhere for 2 hours. Cells were treated with 500 nm-carboxylated PS particles or LPS (1 µg/ml) for 18 hours in triplicates. After incubation, cells were lysed with 1 ml of TRIzol reagent (Thermofisher Scientific) and total RNA extracted from cells using RNeasy Mini Kits (Qiagen, USA) and reverse transcribed to complementary DNA (cDNA) using iSCRIPT cDNA synthesis kit (Bio-Rad, USA) as per the manufacturer’s protocol. The cDNA was analyzed for gene expression using TB Green *Premix Ex Taq* I (Takara Bio Inc., Japan). The RNA expression levels were normalized to the levels of the house-keeping gene, glyceraldehyde 3-phosphate dehydrogenase (*Gapdh*). Sequences of primers used for amplification of human genes are shown in Supplementary Table1.

### Membrane fluidity measurements using Laurdan

RAW cells, 75 × 10^3^ per well, were seeded in a 96-well black flat-bottom polystyrene plate and allowed to adhere overnight. Cells were incubated with 500 nm-carboxylated PS particles at a cell to particle ratio of 1:50 or 1:200 for 2 hours. As a positive control, cells were treated with 66 µM Methyl-β-cyclodextrin (MβCD) (Merck, USA) for 1 hour. After incubation, media was removed, wells washed with 1 X PBS and incubated with 50 µL of incomplete media containing 5 µM Laurdan (Avanti^®^ Polar Lipids, Merck, USA) for 30 min at 37°C. Fluorescence was measured with a microplate reader (Tecan, Switzerland). Emission intensity was acquired at 440 and 490 nm (excitation=385 nm, 5 nm bandpass) at 37 °C. Generalized polarization^50^(GP_laurdan_) was calculated from the emission intensities using the following equation:

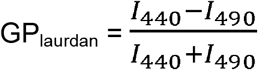

I_440_ and I_490_ represent the fluorescence intensity emitted at 440 nm and 490 nm, respectively.

### Analysis of cellular stiffness using atomic force microscopy (AFM)

RAW cells were seeded at a density of 2 × 10^5^ cells per ml in a 35-mm cell culture dish and allowed to adhere overnight. Particles were added to cells at a ratio of 1:200 for 2 hours to ensure uptake by >95 % cells. Extracellular particles were removed by washing twice with 1 X PBS and cells maintained in DMEM with 10% FBS for AFM measurements. The apparent modulus of elasticity of the cells was measured using an Atomic Force Microscope (XE Bio from Park Systems, Suwon, South Korea) using a V-shaped cantilever (stiffness 0.041 N/m as measured using thermal tuning) with a spherical bead of diameter 5.2 μm made of silicon dioxide (AppNano HYDRA6V-200NG-TL; AppNano, Mountain View, CA). One force-displacement (F-d) curve was obtained per cell for a total of approximately 20 cells in each of the three independent experiments. The force curve corresponding to the approach of the tip towards the substrate was measured, and a constant indentation depth of 200 nm was maintained. Each of these F-d curves was analyzed to obtain the apparent modulus of elasticity of the cell using the Hertzian contact model.^51^

### *In vitro* Bactericidal assays

RAW cells were seeded at a density of 1 × 10^5^ cells per well in a 48-well plate and incubated with 500 nm-carboxylated PS particles, in triplicates, for 18 hours. The cell to particle ratio was 1:50, which was optimized (by testing multiple cells to particle ratios – data not shown) to obtain more than 90% for cells with particles. Simultaneously, *E. coli* K12 MG1655 or *Salmonella enterica* serovar Typhimurium (*S*. Typhimurium) cultures were grown in Luria-Bertani (LB) media at 37 °C till the late log phase (10 hours). The bacterial load was confirmed by plating the cultures on LB-agar plates and enumerating the colony-forming units (CFU). Antibiotic-free DMEM cell culture media was used for further steps in the experiment. After incubating cells with particles, the wells were washed with 1 X PBS to remove non-phagocytosed particles. Bacteria were then added to the cells at the desired multiplicity of infection (MOI), and the plate was centrifuged at 400 rcf for 5 min to allow the bacteria to settle. The plate was incubated at 37°C under 5% CO_2_ for one hour to allow phagocytosis of bacteria by RAW cells. After this, media was removed, wells washed with 1 X PBS, and cells were further incubated in media containing 100 µg/ml gentamicin (HiMedia Laboratories, India) for one hour to kill extracellular bacteria. The viable bacteria inside RAW cells at different time points post-infection were enumerated by lysing the cells with 0.1% Triton X-100 for 10 min and plating the cell lysates on LB-Agar plates to determine bacterial CFU. For all time points beyond 2 hours post-infection, wells were washed with 1 X PBS and cells were maintained in media containing 25 µg/ml of gentamicin.

### *n vivo* killing assays-*E. coli*

C57BL/6 mice (weighing 20 – 30 grams, 8-14-week-old) were injected intraperitoneally 1 × 10^10^ 500 nm-carboxylated PS particles. Control mice were injected with saline. After 2 hours, mice were injected intraperitoneally with ∼5 × 10^7^ *E. coli*. Next, 2- or 6-hours of bacterial injection, mice were euthanized, peritoneal lavage performed with 5 ml of ice-cold PBS-EDTA, and peritoneal exudates collected. The peritoneal exudate cells were stained for live dead using Zombie Aqua Fixable Viability dye (BD Biosciences), fixed with 1.6% paraformaldehyde (PFA), and finally stained with the following antibodies for 20 min at 4 °C: Ly6G (1A8), Ly6C (AL-21), CD11b (M1/70) and F4/80 (T45-2342) and analyzed on a flow cytometer. Further, mice were dissected after peritoneal lavage to collect spleen, kidney, and liver. The organs were mechanically homogenized in 1 X PBS containing 0.1 % Triton X-100 using a pestle. Serial dilutions of each lysate were plated on LB agar plates and incubated overnight at 37 °C. CFU obtained were enumerated to determine the bacterial load for each organ.

### *In vivo* killing assays-*S. epidermidis*

*S. epidermidis*: sGFP expressing plasmid (pTH100-Addgene plasmid #84458)^52^ was transformed into *E. coli* DC10B cells.^53^ Plasmids extracted from DC10B cells were electroporated into *S. epidermidis* (ATCC122293) strain. The cells were plated onto Tryptic soya agar (TSA) plates containing 10 µg/ml chloramphenicol and incubated at 30°C for 24 hours. Fluorescent green colonies were picked for further experiments.

C57BL/6 mice (weighing 20 – 30 grams, 8-14-week-old) were injected intraperitoneally with 1 × 10^9^ 500 nm-carboxylated PS particles. Control mice were injected with saline. After 2 hours, mice were injected intraperitoneally with ∼3 × 10^8^ *S. epidermidis*. After 3- or 24 hours of *S. epidermidis* injection, mice were euthanized, peritoneal lavage performed with 5 ml of ice-cold PBS-EDTA, and peritoneal exudates collected. The peritoneal exudate cells were stained for live dead using Zombie Aqua Fixable Viability dye (BD Biosciences (USA), fixed with 1.6% PFA, and finally stained with the following antibodies for 20 min at 4 °C: CD11b (M1/70), F4/80 (T45-2342) Ly6G (1A8), Ly6C (AL-21), and analyzed on a flow cytometer. Further, mice were dissected after peritoneal lavage to collect spleen, kidney, and liver. The organs were mechanically homogenized in 1 X PBS containing 0.1 % TritonX-100 using a pestle. Serial dilutions of each lysate were plated on Tryptic Soya agar plates and incubated overnight at 37 °C. CFU obtained were enumerated to determine the bacterial load for each organ.

### Expression of activation markers on peritoneal macrophages

C57BL/6 mice were euthanized, and peritoneal lavage was performed with 5 ml of ice-cold PBS-EDTA. Peritoneal exudates were centrifuged at 400 rcf for 5 min, and the cell pellet was resuspended in DMEM complete media. Cells were counted using a hemocytometer. 5 × 10^5^ cells were seeded in each well of a 6-well plate and incubated at 37°C for 2 hours to allow peritoneal macrophages to adhere. Non-adhered cells were removed by washing the wells thrice with 1 X PBS. Fluorescent-labeled 500 nm-carboxylated PS particles were added to cells at a ratio of 1:100 (low) or 1:1000 (high) and incubated for 2 hours. Cells were washed with ice cold PBS-EDTA (three times), scraped and collected in FACS tubes, and stained using Zombie Aqua Fixable Viability dye. Finally, cells were stained with a combination of the following antibodies for 30 mins at 4°C: CD11b (M1/70), CD38 (90), CD54 (3E2), CD62L (MEL-14), CD86 (GL-1), F4/80 (T45-2342) and Ly6C (AL-21) (BD Biosciences). The expression of activation markers was determined using a flow cytometer.

### Statistics

At least three independent experiments were performed (unless explicitly stated differently), and data from biological duplicates of a single independent experiment are reported through a single mean value. For data involving comparisons between 2 groups, Student’s *t*-test or Mann-Whitney test or Welch’s *t*-test were used. For data involving comparisons between multiple groups, one-way ANOVA followed by the Bonferroni *post-hoc* test was used for statistical comparisons.

## Acknowledgements

We thank Monisha for assistance with the atomic force microscope. We acknowledge the support of the staff at the central animal facility and flow cytometry facility, IISc. This work was supported by the DBT/Wellcome Trust India Alliance Fellowship [grant number IA/I/19/1/504265] awarded to SJ. This work was partly supported by a Science and Engineering Board, Department of Science and Technology, Govt. of India [grant number ECR/2016/000629] to SJ. This work was also partly supported by the R. I. Mazumdar young investigator fellowship at the Indian Institute of Science. PS is supported by a research fellowship from University Grants Commission, Govt. of India.

