## Supplementary for "Particle Uptake Driven Phagocytosis in Macrophages and Neutrophils Enhances Bacterial Clearance"



| Item | Page Number |
|---|---|
| Supplementary Figures | 2 – 11 |
| Supplementary Table | 12 |

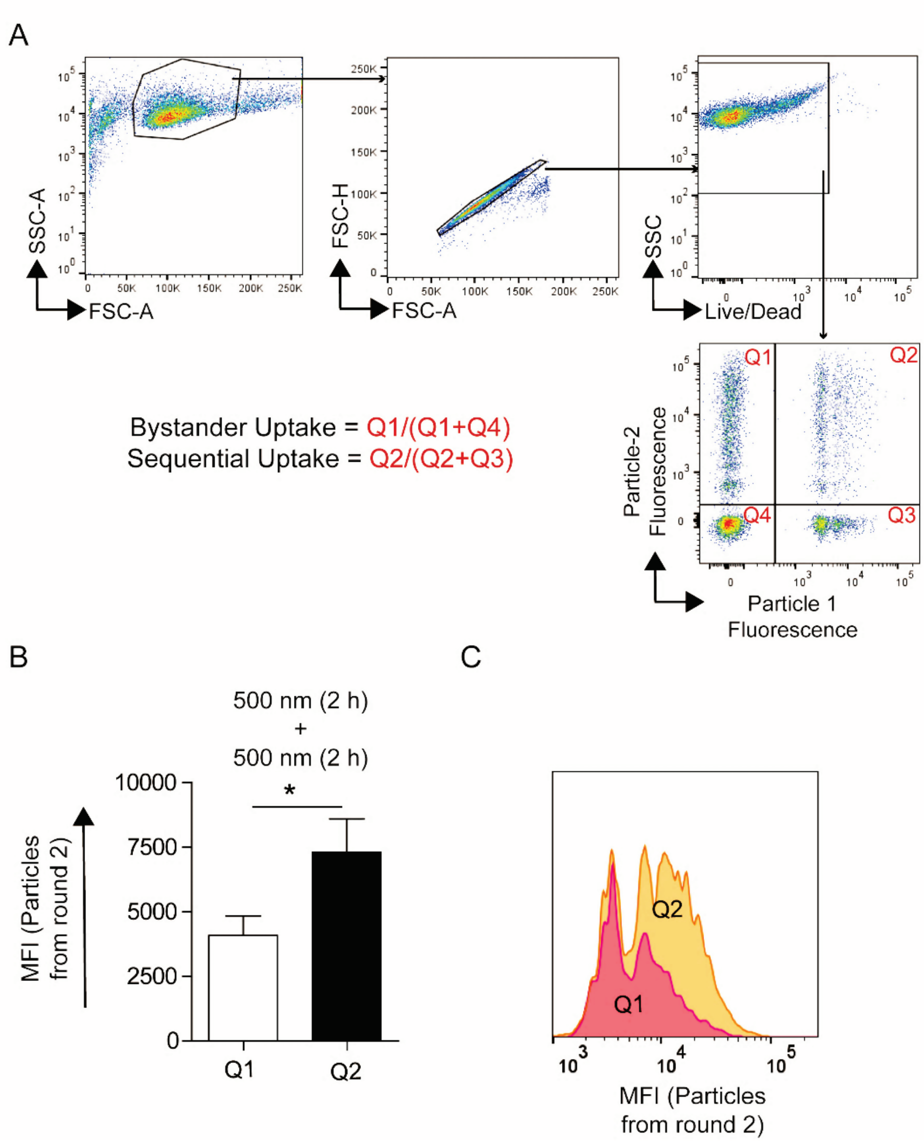


**Supplementary Figure 1: A –** Representative flow cytometry dot plots describing the gating strategy used to quantify bystander and sequential uptake by RAW macrophages (similar strategy was used for all other cell types too). **B** and **C –** Median fluorescence intensity (MFI) as a measure of the number of particles phagocytosed in the second round of phagocytosis by bystander cells (Q1) and cells with 500 nm-carboxylated polystyrene (PS) particles from round 1 (Q2). Data sets are representative of $n \geq 3$. * = $p < 0.05$ calculated using a paired Student's *t*-test.

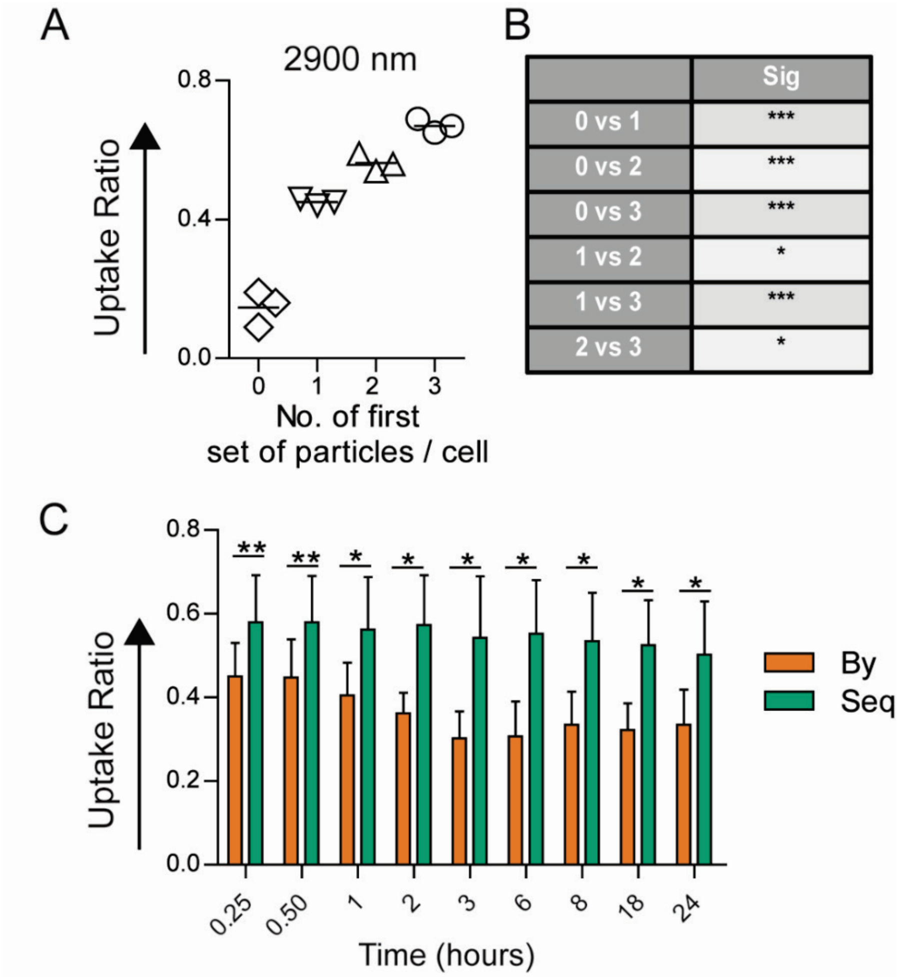


**Supplementary Figure 2: A** – Sequential uptake measured when cells take up different number of particles (2900 nm-carboxylated polystyrene particles added to cells at a cell to particle ratio of 1:2). **B** – Statistical comparison of data shown in **A** (*n* = 3), performed using one-way ANOVA. **C** – Enhanced sequential phagocytosis observed at different incubation times with particles in first round of phagocytosis (t1 in figure 1A). Experiment was performed by addition of 1 µm-carboxylated PS particles and 500 nm-carboxylated PS particles (ratio of 1:50) in a sequential manner. Data sets are representative of $n = 3$. * = $p < 0.05$ and ** = $p < 0.01$ calculated using paired Student's *t*-test.

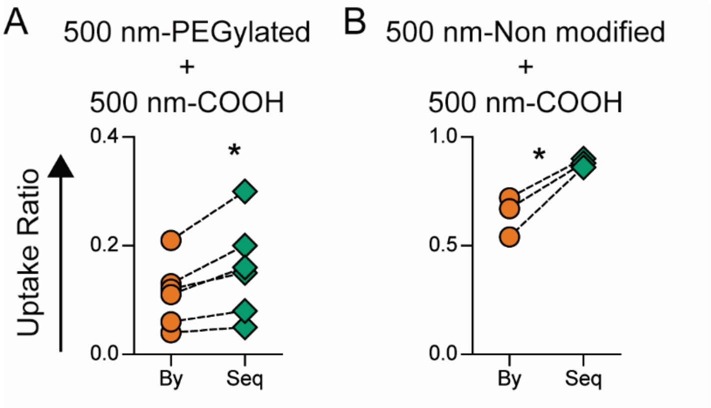


**Supplementary Figure 3:** Enhanced sequential phagocytosis observed with particles having different surface modifications. **A** – Polyethylene glycol (PEG) conjugated 500 nm polystyrene (PS) particles or **B** – 500 nm PS particles with no surface groups were added to RAW cells in first round (cell to particle ratio of 1:50 for 2 hours) followed by addition of 500 nm-carboxylated PS particles (cell to particle ratio of 1:50 for 2 hours) in the second round. Bystander (By) and sequential (Seq) uptake was quantified using flow cytometry. Data sets are representative of $n \geq 3$. * = $p < 0.05$ calculated using paired Student's *t*-test.

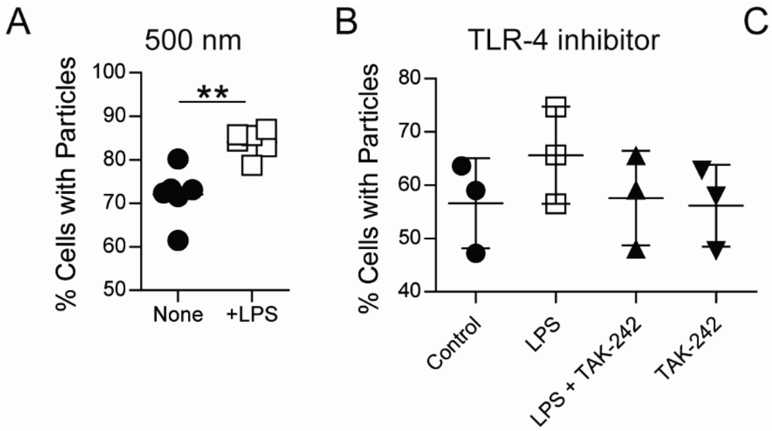


C

| | Sig |
|---|---|
| Control vs LPS | *** |
| Control vs LPS+TAK-242 | ns |
| Control vs TAK-242 | ns |
| LPS vs LPS+TAK-242 | *** |
| LPS vs TAK-242 | *** |
| LPS+TAK-242 vs TAK-242 | ns |

**Supplementary Figure 4: A** – LPS treatment results in increased uptake of particles by RAW macrophages. Cells were cultured with LPS (1 μg/ml) for 12-18 hours, and 500 nm-carboxylated polystyrene (PS) particles added subsequently at a cell to particle ratio of 1:50 for 2 hours. Percentage uptake was determined using a flow cytometer. Data sets are representative of $n = 6$. ** = $p < 0.01$ calculated using Student's *t*-test. **B** – Uptake of 500 nm-carboxylated PS particles, added at a cell to particle ratio of 1:50, measured after treatment with LPS, LPS+TAK-242, or TAK-242 only. LPS treated cells show significantly higher uptake as measured by one-way ANOVA shown in **C**, and TAK-242 inhibits LPS mediated increase in uptake. Data sets are representative of $n = 3$. ** = $p < 0.01$, *** = $p < 0.001$ and ns = non-significant.

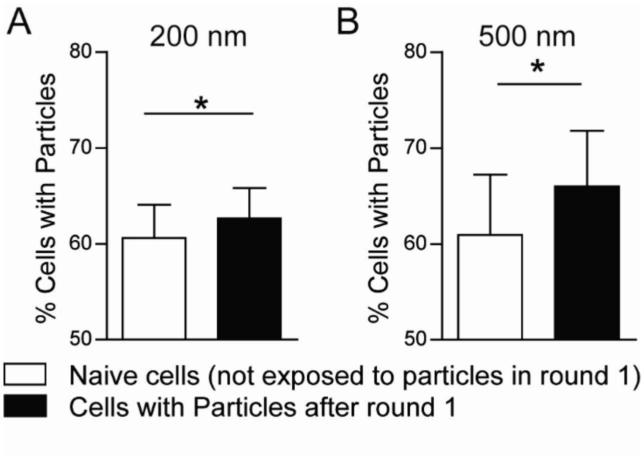


**Supplementary Figure 5: Uptake measure in cells that have particles (experimental group) compared to particle-naïve cells (control group).** Cells were cultured with 200 nm (cell to particle ratio of 1:4000 incubated for 2 hours) or 500 nm (cell to particle ratio of 1:200 incubated for 2 hours) carboxylated PS particles (experimental groups) such that over 95% of cells had particles. In these cells, the ability to take up additional particles was quantified by adding particles made of a different fluorophore (1 µm-carboxylated PS at a cell to particle ratio of 1:10) and compared to naïve cells (not exposed to particles in the first round), which were exposed to the particles in the “second round”. Data are based on $n \geq 3$ independent experiments (each performed in duplicate). * indicates $p < 0.05$ measured using paired Student’s *t*-test.

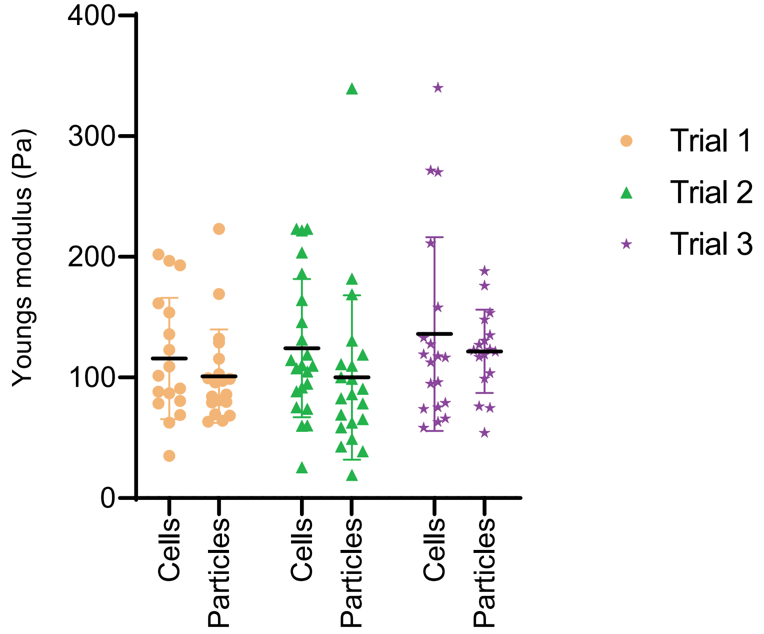


**Supplementary Figure 6:** Data presented in figure 5B represented as independent trials

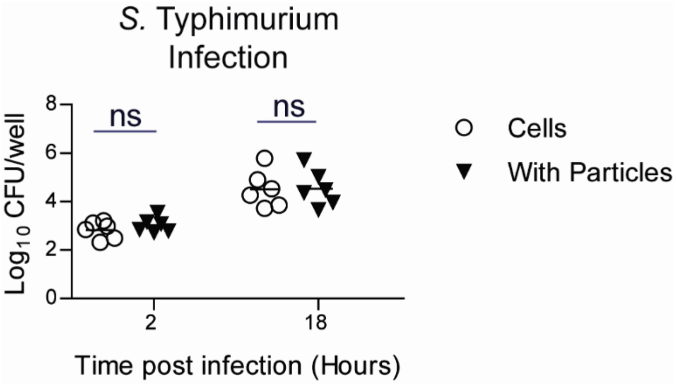


**Supplementary Figure 7: *In vitro* bacterial killing assay: *S.* Typhimurium**
Uptake and intracellular replication of *S. Typhimurium* (MOI 5-50) inside RAW macrophages following culture of RAW cells with 500 nm-carboxylated polystyrene particles (or control cells without particles). Intracellular bacterial numbers were determined using gentamicin protection assay at various times post-infection. Enumeration of intracellular bacteria measured as colony forming units (CFU) obtained by plating cell lysate at specified times post infection. Data sets are representative of $n$ = 6 independent experiments. ns indicates non-significant calculated using Student's $t$-test.

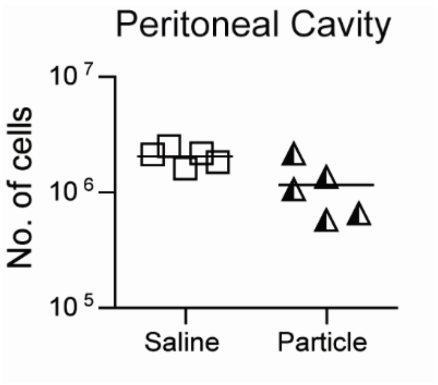


**Supplementary Figure 8:** Total cell counts in peritoneal cavity of mice 2 hours following intraperitoneal injection of $10^{10}$ 500 nm-carboxylated polystyrene particles compared to saline injected mice. Each data point corresponds to one mouse. Mann-Whitney test was performed for statistical comparison.

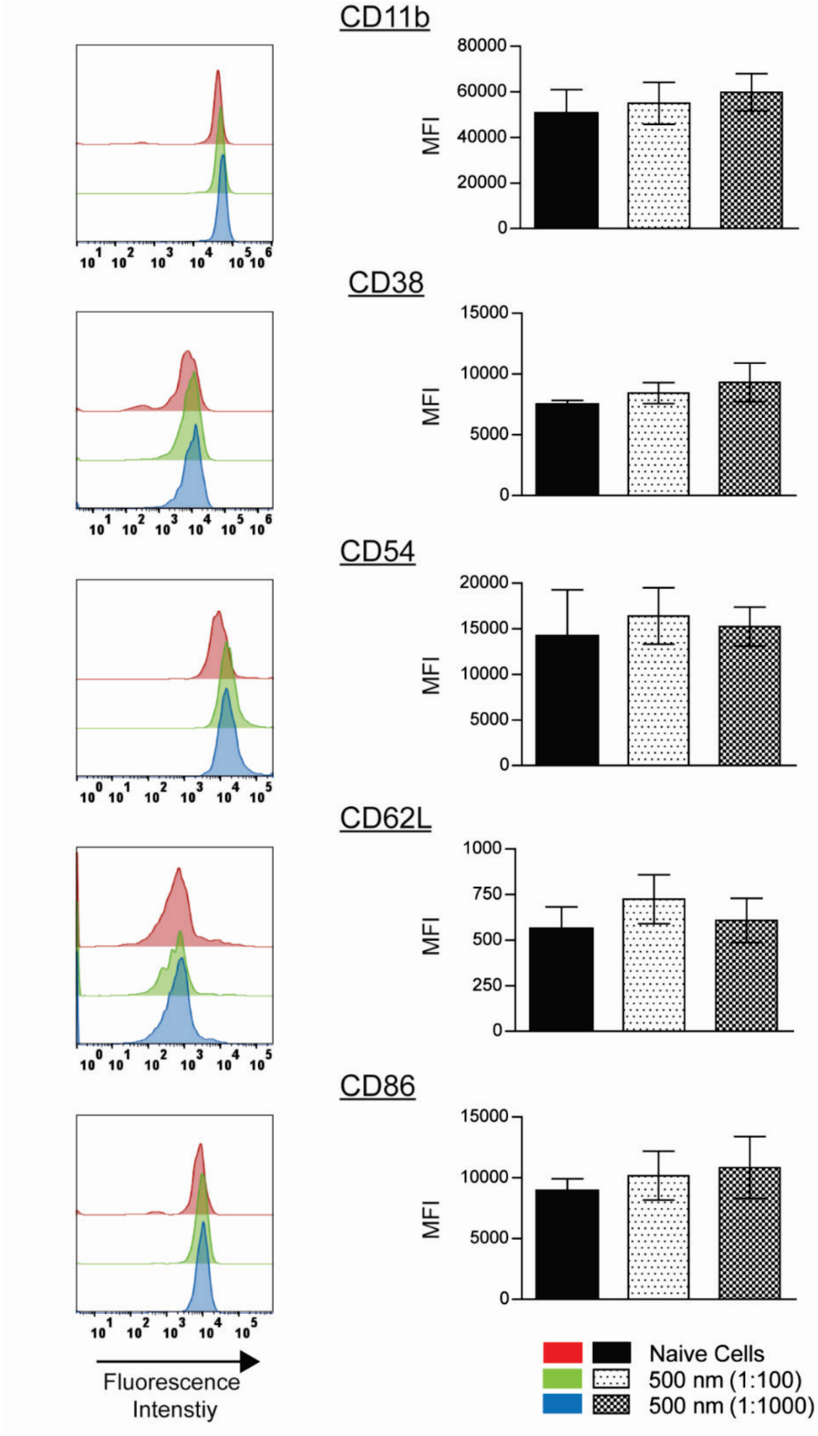


**Supplementary Figure 9:** Expression of surface proteins CD11b, CD38, CD54, CD62L and CD86 on mouse peritoneal macrophages incubated with 500 nm-carboxylated polystyrene particles for 2 hours and compared to naïve cells that were not exposed to particles. Data are based on *n* = 3 independent experiments (each performed in duplicate). One-way ANOVA showed no statistical differences between the groups

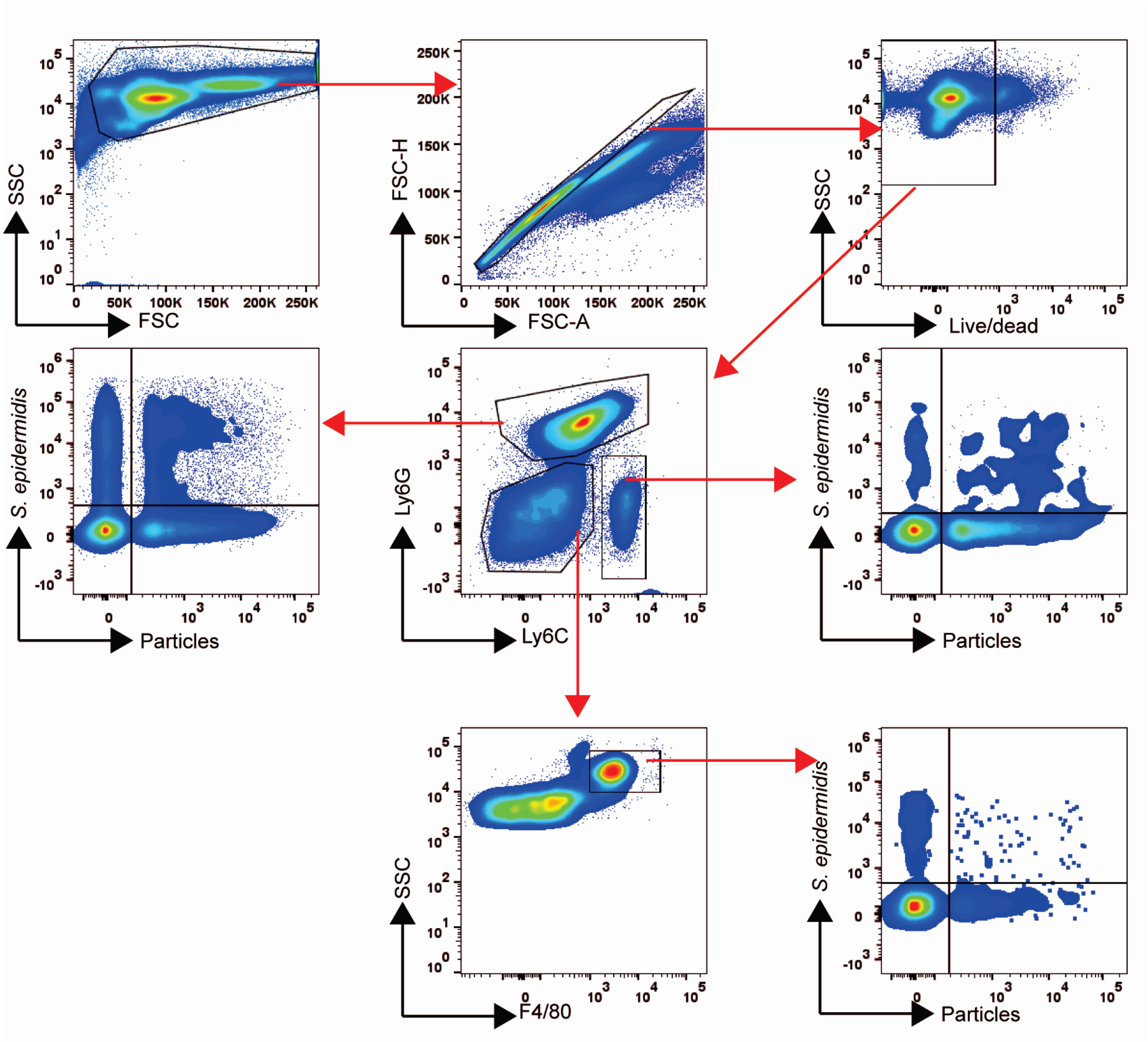


**Supplementary figure 10: Gating strategy used to determine bystander and sequential uptake of *S. epidermidis in vivo.*** Representative flow cytometry dot plots describing the gating scheme used for *in vivo* bacterial killing assays with *S. epidermidis*. Ly6G is a marker for neutrophils, Ly6C is a marker for monocytes, and F4/80 expressing cells that do not express Ly6G and Ly6C are macrophages. SSC indicates side scatter and FSC indicates forward scatter.

**Supplementary Table 1: List of Primers and primer sequence used**

| Gene | Forward Primer | Reverse Primer |
|---|---|---|
| ***Gapdh*** | 5’-AGGTCGGTGTGAACGGATTTG-3’ | 5’-GGGGTCGTTGATGGCAACA-3’ |
| ***IL-1β*** | 5’-CAACCAACAAGTGATATTCTCCATG-3’ | 5’-GATCCACACTCTCCAGCTGCA-3’ |
| ***IL- 6*** | 5’-TACCACTTCACAAGTCGGAGGC-3’ | 5’-CTGCAAGTGCATCATCGTTGTTC-3’ |
| ***Tnf-α*** | 5’-GGTGCCTATGTCTCAGCCTCTT-3’ | 5’-GCCATAGAACTGATGAGAGGGAG-3’ |
